# The domain-within-domain packing of euchromatin can be described as multiplicative cascades

**DOI:** 10.1101/2020.10.20.346882

**Authors:** Amra Noa, Hui-Shun Kuan, Vera Aschmann, Vasily Zaburdaev, Lennart Hilbert

## Abstract

The genome is packed into the cell nucleus in the form of chromatin. Biochemical approaches have revealed that chromatin is packed within domains, which group into larger domains, and so forth. Such domain-within-domain packing, also called hierarchical packing, is equally visible in super-resolution microscopy images of large-scale chromatin organization. While previous work has suggested that chromatin is partitioned into distinct domains via microphase separation, it is unclear how these domains organize into a hierarchical packing. A particular challenge is to find an image analysis approach that fully incorporates such hierarchical packing, so that hypothetical governing mechanisms of euchromatin packing can be compared against the results of such an analysis. Here, we obtain 3D STED super-resolution images from pluripotent zebrafish embryos labeled with improved DNA fluorescence stains, and demonstrate how the hierarchical packing of euchromatin in these images can be described as multiplicative cascades. Multiplicative cascades are an established theoretical concept to describe the placement of ever-smaller structures within bigger structures. Importantly, these cascades can generate artificial image data by applying a single rule again and again, and can be fully specified using only four parameters. Here, we show how the typical patterns of euchromatin organization are reflected in the values of these four parameters. In particular, we can pinpoint the values required to mimic a microphase-separated configuration of euchromatin. We suspect that the concept of multiplicative cascades can also be applied to images of other types of chromatin. In particular, cascade parameters could serve as test quantities to assess whether microphase separation or other theoretical models accurately reproduce the hierarchical packing of chromatin.

**SIGNIFICANCE:** DNA is stored inside the cell nucleus in the form of chromatin. Chromatin exhibits a striking three-dimensional organization, where small domains group into larger domains, which again group into larger domains, and so forth. While this hierarchical domain-within-domain organization is obvious from microscopy images, it is still not entirely clear how it is established, or how it should be properly characterized. Here, we demonstrate that multiplicative cascades – a concept from theoretical physics used to characterize for example cloud patterns, galaxy locations, or soil patterns – are also ideally suited to describe the domain-within-domain organization of chromatin. This description is rather simple, using only four numbers, and can thus facilitate testing of competing theories for the domain-within-domain organization of chromatin.

## INTRODUCTION

The packing of a several meters long genome into a cell nucleus of only a few micrometers diameter is often referenced as a remarkable phenomenon. Another striking observation is that, across all investigated length scales, this packing follows an apparent hierarchical organization. Specifically, light and electron microscopy, and more recently biochemical methods in combination with modern DNA sequencing technology, have revealed how small domains of chromatin come together to form larger domains, which in turn come together to form yet bigger domains, and so forth (1–3). Microscopy approaches have shown hierarchical organization in the chromatin fibre at ≈ 10 nm (4, 5), amongst chromatin domains at at the scale of ≈ 100 nm (6–9), and also for compartments and territories on the micrometer scale (10). In the context of biochemical studies, which investigate genome organization via the point-to-point contacts of different positions on chromosomes, the observation of such hierarchical packing has been explained by the fractal globule model (11–13), and more recently by the loop-extrusion model (14, 15). It is, however, still poorly understood how the domain-within-domain packing as visualized in microscopy images is established.

We have recently addressed the internal patterning of euchromatin in relation to transcription activity, and proposed that a fine-grained pattern of euchromatin domains is established and maintained in line with a microphase-separated state of a microemulsion (16). During our work with super-resolution images of euchromatin organization, it has become increasingly apparent that euchromatin also exhibits a domain-within-domain structure. One difficulty in our work was that, to our knowledge, no analysis approaches were in common use that would pay full attention to such a structure. However, for the understanding of euchromatin organization as well as potential underlying microphase separation mechanisms, such an analysis would be highly beneficial. Here, we visualize euchromatin organization in zebrafish embryonic cells by 3D STimulatd Emission Depletion (STED) super-resolution microscopy using improved DNA stains, and demonstrate how the euchromatin organization in the images can be described in terms of multiplicative cascades. Multiplicative cascades are a well-established concept from theoretical physics, used to describe how patterns are formed by splitting a large structure into smaller and smaller sub-structures, while consistently adhering to one and the same rule how to execute this splitting. Importantly, a wide variety of complex patterns can be generated based on only four numbers that fully specify the cascade process that leads to the final pattern. Such multiplicative cascades seem ideally suited as a simple description of the hierarchical organization of chromatin domains, as shown here for the example of euchromatin that is organized in line with a microphase separation scenario.

## MATERIALS AND METHODS

### Collection and staining of zebrafish embryos

Late blastula zebrafish embryos were collected and stained as described previously (17). Briefly, embryos of wild type zebrafish (AB, sourced from the Zebrafish International Resource Center and maintained at the European Zebrafish Resource Center) were obtained by spontaneous mating and dechorionated by pronase treatment. Embryos were collected and fixed (2% formaldehyde, 0.2% Tween-20) at the sphere stage of development, and subsequently permeabilized (0.5% Triton X-100, 15 minutes at room temperature). Mounting media (Vectashield H-1000, Vector Laboratories, Burlingame, CA, USA; Glycerol; TDE-O, Abberior, Göttingen, Germany) and DNA stains were mixed and applied to the samples on the day of microscopy, as fluorescence intensity decayed noticeably overnight. Fluorescence stains were used at 10 *μ*M (SiR-DNA, Spirochrome, Stein am Rhein, Switzerland) or 10x recommended dilution (SPY650-DNA, SPY595-DNA, both Spirochrome, molarity not indicated by manufacturer). Embryos were cleaned of yolk particles and placed under #1.5 selected coverslips with spacers.

### Microscopy

Microscopy images were recorded with a Leica TCS SP8 STED microscope (Leica Microsystems, Wetzlar, Germany) equipped with a wavelength-adjustable white light laser, a 775 nm depletion line, and time-gated HyD detectors, using a motorized-correction 93x NA 1.30 glycerol objective (HC PL APO 93X/1.30 GLYC motCORR). The STED line was used in 100% 3D depletion mode throughout. Within a given experiment and dataset, all illumination and acquisition settings were maintained unchanged, so as to obtain quantitatively comparable images. For each acquired image or 3D stack, the correction collar setting was optimized by adjustment for maximal image intensity (18).

### Image analysis

Staining intensities were quantified using Cell Profiler. Specifically, nuclei were segmented based on the DNA fluorescence intensity using a global Otsu threshold algorithm. The nuclear segmentation mask was dilated to obtain cytoplasmic mask.

Image contrast (*C*_*DN A*_) and correlation length (*L*_*corr*_) values were calculated in Matlab. *C*_*DN A*_ was calculated as the coefficient of variation of intensity values (microscopy data) or concentration values (generated concentration profiles), dividing the standard deviation by the mean. In the generated data, a correction factor of 0.45 was applied the *C*_*DN A*_ values, to adjust for fluorescence background in the microscopy data (16). *L*_*corr*_ = *C*^1/4^ was determined by fitting the function

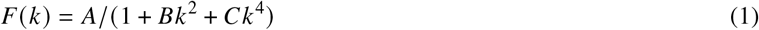

to the structure factor calculated from the image intensities. The fitting function relates to the common physical structure factor *S*(*k*) in the following relation (19):

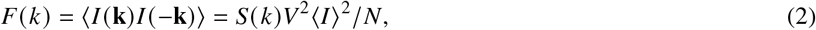

where *I*(**k**) is the two-dimensional spatial Fourier transform of the image intensity values (*I*_*ij*_), 〈…〉 denotes the radial average, *i* and *j* the 2D indices in the analyzed plane of the image stack, so that 〈*I*〉 is the average intensity. *V* is the system volume, and *N* denotes the microscopic number of particles. Here, the linear assumption between the image intensity and the particle density *N*/*V* is considered.

The fitting function *F*(*k*) is taken from microemulsion theory (20). From the order parameter *Ψ* expansion, the free energy density is written as *f* = *aΨ*^2^ + *c*_1_ (∇*Ψ*)^2^ + *c*_2_ (∇^2^*Ψ*)^2^, where the corresponding structure factor *S*(*k*) is proportional to (*a* + *c*_1_*k*^2^ + *c*_2_*k*^4^)^−1^ and it is also proportional to *F*(*k*) in our system via equation (2).

### Multi-fractal analysis

The multi-fractal features of the images are analyzed with the box counting method (21, 22). The image is separated into *N*(*L*) sub-images with the size *L* × *L*, where *L* has the unit of pixels. The normalized intensity of the sub-image *i* is *P*_*i*_(*L*) = *I*_*i*_(*L*)/*I*_*t*_, where *I*_*i*_(*L*) is the intensity in the sub-image *i* and *I*_*t*_ = Σ_*i*_ *I*_*i*_(*L*). The generalized fractal dimension is defined as

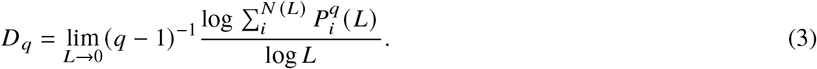

The multi-fractal spectrum, or the so called *f* –*α* curve, is defined by

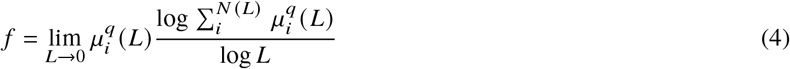

with

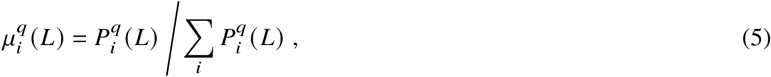

and

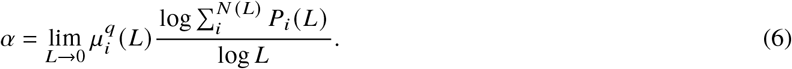

To approximate limit expressions, we performed linear fits over different *L* values.

### Generation of concentration profiles by multiplicative cascades

Cascades were carried out based on the four probabilities *P*_1_, *P*_2_, *P*_3_, and *P*_4_ as a parameter set (0 ≤ *P*_1_, *P*_2_, *P*_3_, *P*_4_ ≤ 1). The cascades were initialized in a 2-by-2 pixel matrix, and pixels were recursively sub-partitioned assigning the four probability values randomly to the four new pixels (Fig. 3B). The recursion was repeated till an image of 256-by-256 pixel size (Fig. 3) or 128-by-128 pixel size (Fig. 4) was obtained. The concentration in this pixel was then calculated as the product of all probabilities assigned to this pixel during the recursion process. A pixel size of 30 nm was assigned, closely matching the pixel size in the microscopy data. Lastly, generated microscopy images were obtained by applying a Gaussian blur with *σ* = 50 nm that approximates the limited resolution of the microscopy images.

### Estimation of multiplicative cascade parameters

An ensemble of 10,000 concentration profiles were generated, using *P*_1_, *P*_2_, *P*_3_, *P*_4_ values randomly drawn from a uniform distribution. For each generated image and each microscopy image, the ‘’mass” present at a given pixel in that image was calculated, *m*_*i*_ = *I*_*i*_/Σ_*i*_ *I*_*i*_. Here *I*_*i*_ is the intensity at the pixel *i*. The mismatch between the mass distribution of two images was then calculated with the error metric

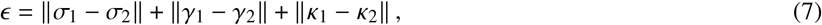

where *σ*_1,2_ are the variance values, *γ*_1,2_ the skewness values, and *κ*_1,2_ the kurtosis values of the mass distribution *m*_i_ of the images 1 and 2. For a given “target” microscopy image, the “top-100” generated images were then chosen based on the lowest values of *ε*. From these top-100 generated images, a single image could further be chosen by the smallest residual error between the radial pair-wise correlation functions of the generated images and the target image (Fig. 4).

## RESULTS

To assess large-scale euchromatin organization, we recorded 3D STED super-resolution microscopy data from nuclei of pluripotent cells in zebrafish embryos. At the pluripotent stage of development (late blastula), these cells exhibit a euchromatin-only nuclear organization and are distributed throughout the different phases of the cell cycle (16, 23, 24). Both aspects have proven beneficial for the study of large-scale euchromatin organization by STED super-resolution microscopy in our previous work (16, 17). To further improve image quality, we assessed the performance of STED-capable fluorogenic DNA stains that recently became commercially available. When we tested the established stain SiR-Hoechst (commercialized as SiR-DNA) in fixed embryos, we found a strong influence of the mounting medium (Fig. 1A,B), as expected from our previous results (17). Vectashield almost completely suppressed fluorescence, glycerol allowed for clearly detectable fluorescence, thiodiethanol (TDE) resulted in almost 10-fold higher fluorescence intensity than glycerol. SiR-DNA in TDE, however, exhibited high levels of cytoplasmic signal, leaving us with reservations towards this stain-media combination. The newly available stain SPY650-DNA also gave high intensities in TDE, with lower cytoplasmic signal. However, when only the STED laser was used, still clearly detectable signal was present, suggesting an unfavorable reexcitation of this stain by the STED laser. Another newly available stain, SPY595-DNA, displayed high levels of nuclear signal in TDE, with the comparatively lowest cytoplasmic and reexcitation signal. To assess the overall performance of this stain-medium combination, we recorded full volumetric stacks of nuclei with the STED line in full 3D depletion mode. In these stacks, no apparent bleaching occurred and distinct chromatin domains could be seen both in lateral (XY) as well as axial (Z) direction (Fig. 1C,D). Edge and aberration artifacts that are typical for 3D STED in thick samples could be mostly avoided by careful correction collar adjustment. Considering the preferable evaluation of the SPY595-DNA stain, we proceeded to assess euchromatin organization using this stain in combination with TDE as a mounting medium.

**Figure 1:**
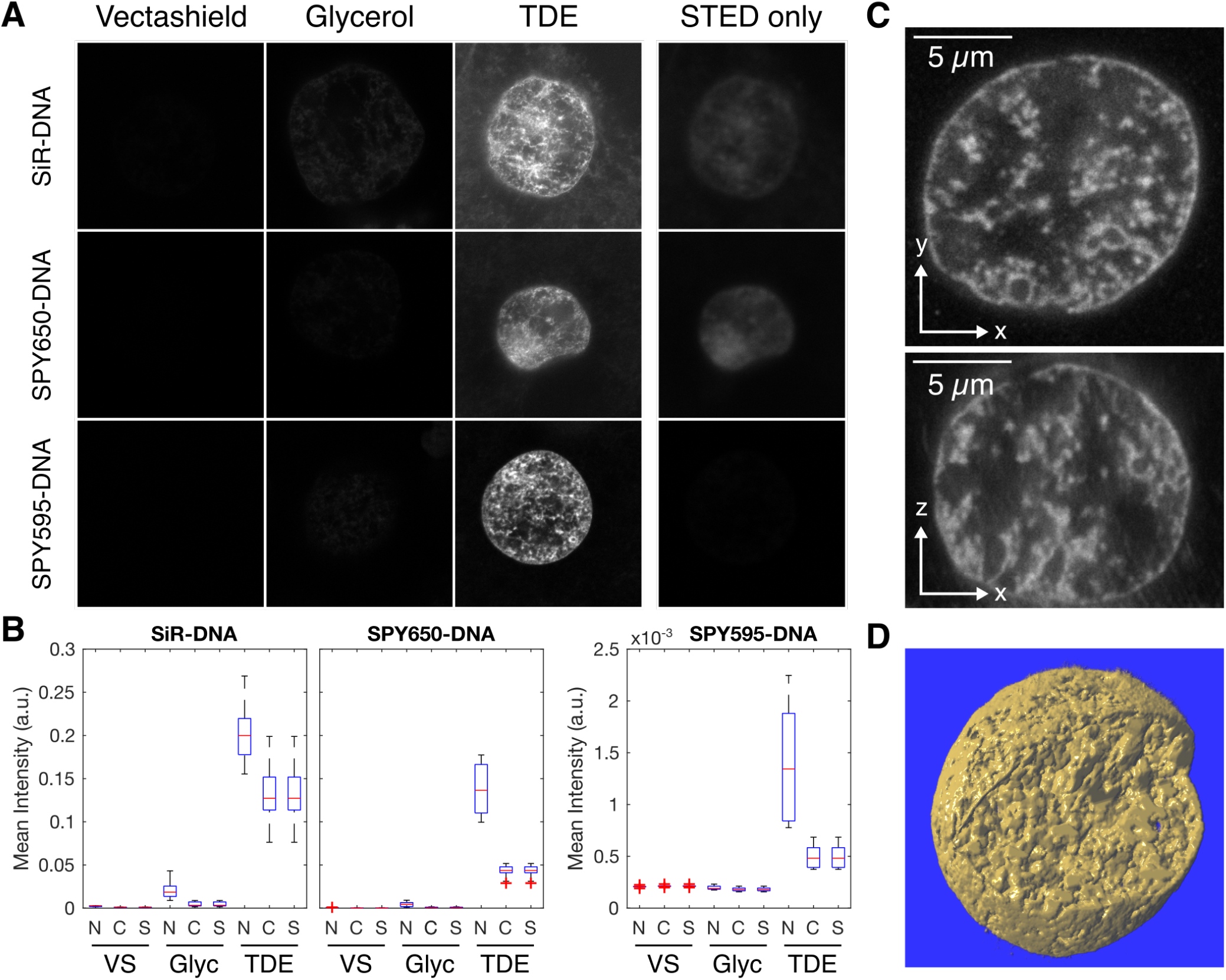
Improved STED DNA stains allow 3D imaging of entire nuclei and reduction of undesirable signal. A) Comparison of STED microscopy signal in an older and two improved commercially available DNA stains. All stains were applied in the mounting media Vectashield, glycerol, and thiodiethanol (TDE). SiR-DNA and SPY650-DNA were excited with 652 nm light, and detected in the window from 670 to 740 nm. SPY595-DNA was excited with 599 nm light, an detected in the window from 510 to 640 nm. No gating was applied in detection. In the STED only condition, only the STED laser but no excitation laser was used on the TDE samples. Images were recorded from oblong stage embryos. B) Quantification of nuclear (N), cytoplasmic (C), and STED only (S) intensity for the different stain-media combinations. C) XY and XZ slices through a full 3D stack of a nucleus in a SPY595-DNA stained sphere stage embryo. D) 3D rendering of data shown in panel C, one quarter of the image stack was cut away to reveal the inside of the nucleus (Volume Viewer ImageJ plugin was used).

We then assessed what different types of euchromatin organization can be found in the different nuclei in the embryos. An overview scan revealed a diverse range of chromatin organization patterns (Fig. 2A). After excluding nuclei of cells preparing for, undergoing, or exiting division, we found three typical patterns of organization: smooth, intermediate, and coarse (Fig. 2A). Based on our previous work, these typical euchromatin patterns can be related to different configurations of a polymer suspension (16). The smooth pattern corresponds to a polymer suspension in a mixed configuration, which is found early in the cell cycle when transcription has not yet recommenced. The intermediate pattern corresponds to a microphase-separated suspension, where euchromatin segregates from RNA-rich regions of the nucleus but remains dispersed into small domains by high levels of transcription. The coarse pattern corresponds to an approach towards a fully phase-separated configuration, which occurs in cells with lower transcription levels. To more systematically assess these types of euchromatin organization, we quantified how prominent the patterns formed by euchromatin were (image contrast, *C*_*DN A*_) and what the characteristic distance is over which euchromatin patterns are formed (correlation length, *L*_*corr*_) (Fig. 2B). The analyzed example images suggest an inverse relationship, where as *C*_*DN A*_ increases, *L*_*corr*_ decreases. Indeed, an analysis over a wider set of nuclei confirms this inverse relationship (Fig. 2C). It appears that the different nuclei form a continuum of euchromatin organization patterns, and the smooth, intermediate, and coarse organization patterns are placed at different positions within this continuum. Such a continuum is in line with previous work, where it was seen that chromatin organization is established progressively after cell division (16, 25). The observed inverse relationship suggests that the progressive establishment of distinct euchromatin domains (increasing *C*_*DN A*_) is correlated with more fine-grained structuring (decreasing *L*_*corr*_). Taken together, our analysis reveals a characteristic inverse relationship of image contrast and correlation length in our ensemble of nuclei. A packing process that captures the observed euchromatin organization should reproduce this relationship.

**Figure 2:**
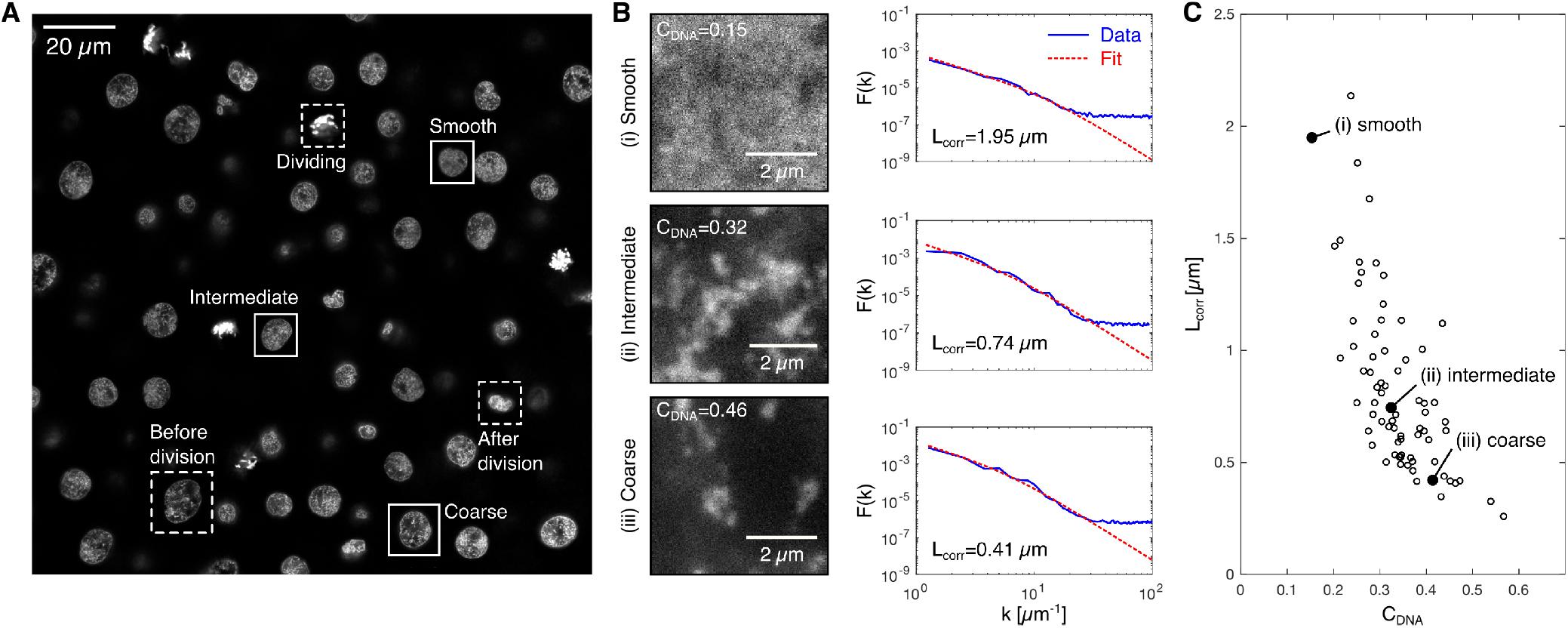
Cell nuclei in the zebrafish embryo display a continuum of euchromatin organization patterns. A) Overview of different chromatin organization patterns in nuclei of a sphere stage zebrafish embryo. Examples of different euchromatin organization patterns in interphase nuclei indicated as solid boxes; examples of nuclei before, during, and after cell division indicated as dashed boxes. Excitation laser 599 nm, detection window 609 to 740 nm, detection gated to within 1.5 to 9 ns after the excitation pulse. B) Representative XY views of image stacks from within nuclei in the smooth, the intermediate, and the coarse euchromatin state. Also shown are image contrast values (*C*_*DN A*_), image-based *F*(*k*) curves, and the fitted *F*(*k*) function used to obtain the correlation length (*L*_*corr*_). C) Correlation length vs. image contrast plotted for 78 nuclei, image data obtained from four different embryos.

The observed range of euchromatin organization patterns points towards a packing process that not only results a domain-within-domain organization, but also allows for a wide range of different patterns. One readily observable natural phenomenon that exhibits (i) a wide range of patterns and (ii) placement of domains within increasingly larger domains, are clouds (Fig. 3A). Due to the observed placement of ever-smaller replicas of domains into larger domains (“self-similarity”), clouds can be related to fractal sets, in particular to multi-fractals (22, 26, 27). Multi-fractals allow for the recognition of self-similarity in systems with stronger spatial inhomogeneity than encountered in conventional fractal sets. When we analyzed our microscopy data, we indeed found that the euchromatin intensity distributions showed properties typical of multi-fractals rather than conventionial fractal sets (SI Fig. 5) (21, 28). Aiming to not only characterize but reproduce the observed intensity distributions, we identified a generative process that is often used to produce multi-fractal patterns, multiplicative cascades (22, 29). Multiplicative cascades generate density profiles by splitting a square space into ever-smaller subsquares, while repeating one and the same splitting rule at every repetition (Fig. 3B). Note that the splitting rule is stochastic and evaluated using a random number generator as part of the image generation process, so that every generated image is different while nevertheless adhering to the same splitting rule. To test the applicability of multiplicative cascades, we generated an ensemble of density distributions using the multiplicative cascade approach. It was previously suggested that the data collection process should be closely replicated when multiplicative cascades are used to reproduce experimental data (30), so we took into consideration the typical image size, physical pixel size, and effective resolution of our microscope. The resulting density distributions contained cases that, leaving aside the expected traces of the square-based cascade procedure, visually resembled the euchromatin organization patterns observed by microscopy (Fig. 3C). Image contrast and correlation length values of these examples were also comparable to the values obtained for the smooth, intermediate, and coarse patterns in our microscopy data (Fig. 3C). When assessing the whole ensemble of generated patterns, the same inverse relationship of *C*_*DN A*_ and *L*_*corr*_ as in our experimental data could be seen (Fig. 3D). Note, however, that the generated patterns spanned a distinctly wider range of values. This comparison of generated patterns to our microscopy data suggests that multiplicative cascades can indeed reproduce key features of the large-scale organization of euchromatin.

**Figure 3:**
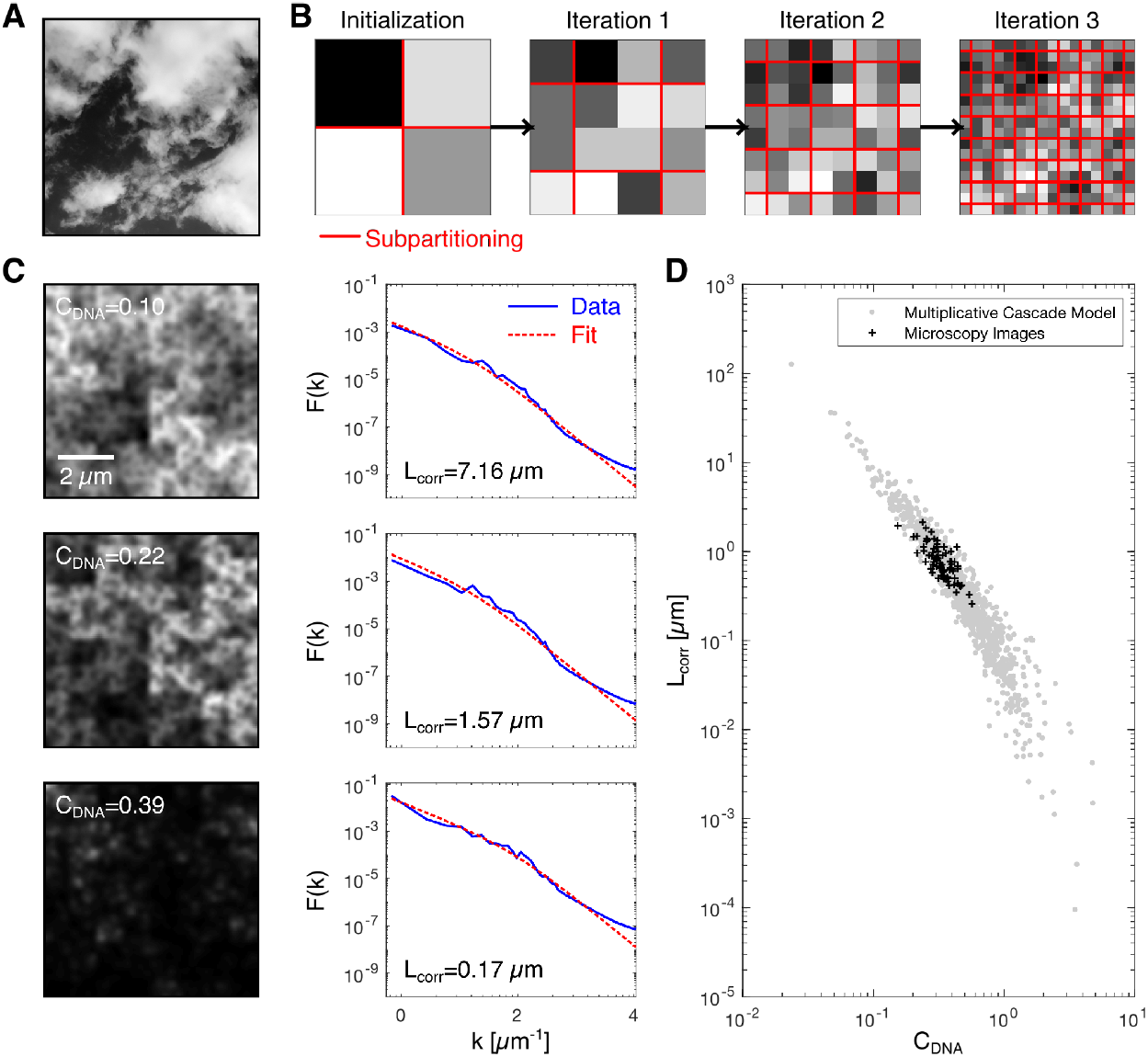
Images generated using multiplicative cascades show similarities with microscopy images of euchromatin. A) Example photograph of a cloud. B) Illustration of the image generation based on multiplicative cascades. In the initial 2-by-2 pixel image, the probabilities *P*_1_, *P*_2_, *P*_3_, *P*_4_ are randomly assigned. During every further iteration, every pixel gets partitioned into four subpixels. As part of this subpartitioning, each pixel is multiplied with one of the probability values *P*_1_, *P*_2_, *P*_3_, *P*_4_, which are randomly allocated. C) Examples of concentration profiles generated by multiplicative cascades, which correspond to the smooth, intermediate, and coarse patterns of euchromatin organization. Calculation of *F*(*k*) curves and extraction of *L*_*corr*_ were carried out identically to microscopy data. D) A set of multiplicative cascades with randomly assigned parameters was executed to allow the comparison of *C*_*DN A*_ and *L*_*corr*_ values against experimental results.

**Figure 4:**
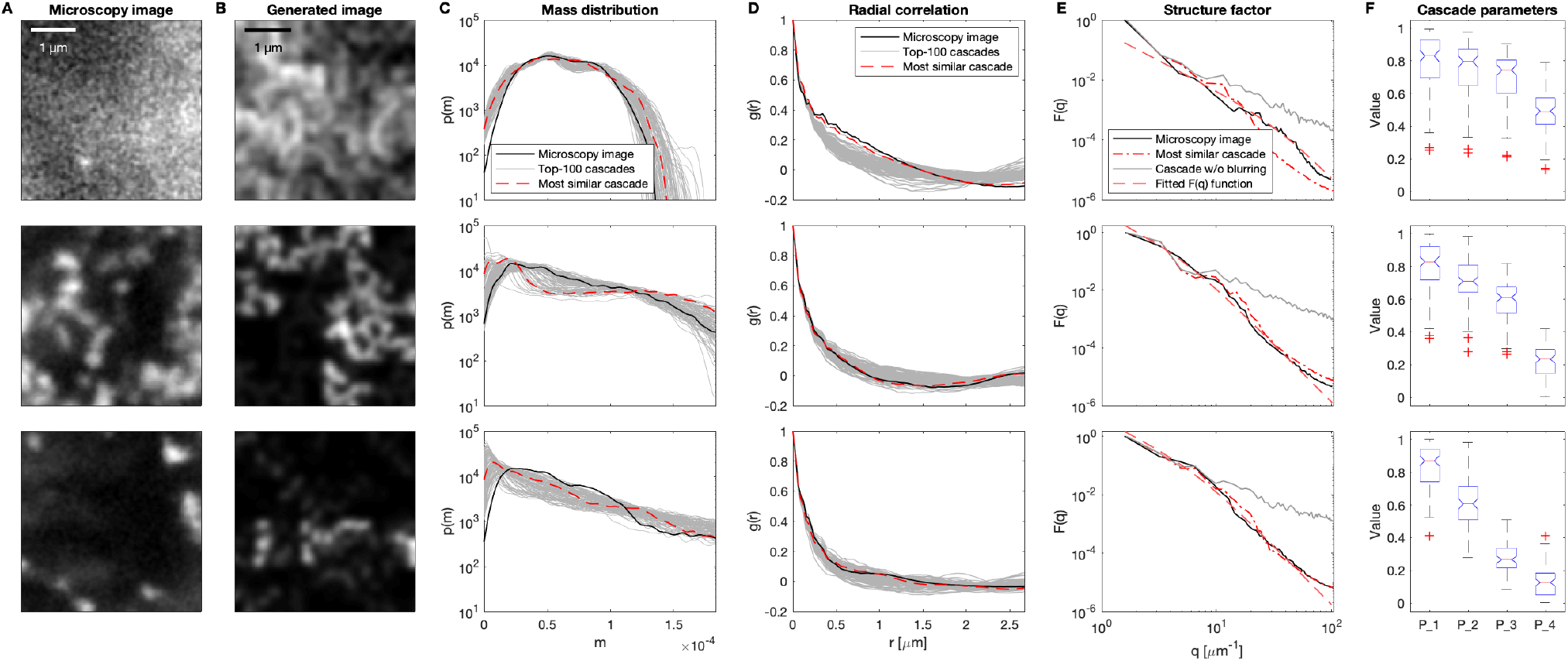
Multiplicative cascades can be empirically adjusted to reproduce euchromatin patterns observed in microscopy images. A) Example microscopy images that were used as targets in the adjustment of multiplicative cascades. Intensities are scaled to the range of each image. B) Generated microscopy images with highest similarity to the microscopy images. C) Mass distributions of the microscopy images, of the most similar generated images, and of the top-100 cascades with highest similarity of mass distribution functions. The top-100 cascades were extracted from an ensemble of 10,000 proposed cascades with randomly assigned parameters. D) Pair-wise radial correlation functions for the microscopy images, the most similar generated images, and the top-100 cascades. E) *F*(*k*) curves for the microscopy data, fit of the microemulsion *F*(*k*) function, *F*(*k*) curve of the raw concentration profiles from the most similar multiplicative cascades, and from the blurred concentration profile from that cascade. F) Boxplots of the cascade parameters of the top-100 cascades.

Now that we established multiplicative cascades as a process that can generate domain-within-domain patterns, we would like to know how the different patterns of euchromatin organization are represented by cascades with specific properties. A given multiplicative cascade process is described by four parameters, *P*_1_, *P*_2_, *P*_3_, *P*_4_. These parameters are the four probabilities that are used during each of the four-way partitioning steps that make up the multiplicative cascade. To fit these parameters to a given target image, we generated an ensemble of 10,000 candidate density profiles with randomly assigned probability parameters. We compared intensity distributions of the generated images resulting from these cascades with the intensity distributions of the target image (Fig. 4A,B). To this end, the “top-100” generated images with mass distribution most similar to the target image were chosen (Fig. 4C). Without further fitting, the “top-100” generated images also agreed well with the radial correlation function of a given target image (Fig. 4D). Out of these “top-100”, the one with the best fitting correlation function was chosen for further investigation.

The key property of multiplicative cascades is that the same cascading rule is applied across all length scales. This is often referred to as scale-free organization. Such scale-free organization seems to be in contradiction with the possibility to quantify a specific correlation length (Fig. 2B). We suspected that the finite resolution of our microscope in combination with the specific patterns of euchromatin organization might be responsible for the observation of an apparent correlation length. In our images generated from multiplicative cascades we recovered scale-free behavior when we left out the blurring step that approximates the finite microscope resolution (Fig. 4E). When we did apply this blurring step, we saw features indicative of a defined length scale (Fig. 4E). This analysis supports a scenario where a multiplicative cascade process, which is inherently scale-free, can reproduce euchromatin organization patterns when the finite microscope resolution is considered.

Given that the fitted cascades seem to represent the experimental data well, we would now like to interpret the parameter values *P*_1_, *P*_2_, *P*_3_, *P*_4_ associated with the different euchromatin patterns. For all euchromatin patterns, a gradual decrease of the probability values starting from *P*_1_ via *P*_2_ and *P*_3_ towards *P*_4_ can be seen (Fig. 4F). Based on the fitting to representative microscopy images, this decrease of the parameter values is the more pronounced the more compacted the euchromatin is in the target images (Fig. 4F). When we further assessed our entire data set of 78 nuclei, this trend was confirmed as statistically significant (SI Fig. 6). Lastly, we paid specific attention to the case of intermediate euchromatin compaction, which corresponds to the microphase-separated state of euchromatin organization. In this case, a clear reduction of *P*_2_, *P*_3_, *P*_4_ was seen. However, this reduction was not as pronounced as for the coarse euchromatin configuration, which is associated with an approach towards full phase separation. We conclude that, indeed, specific patterns of euchromatin organization can be represented by multiplicative cascades with particular parameter sets. The microphase-separated configuration of euchromatin is represented by cascades with a distinct but not maximally pronounced decrease of the parameter values *P*_2_, *P*_3_ and *P*_4_.

## DISCUSSION

We demonstrated how the large-scale euchromatin organization in super-resolution microscopy images can be described in terms of the parameters of a multiplicative cascade. Multiplicative cascades are processes that are not defined at a specific length scale or resolution, but rather describe how smaller and smaller parts of a given structure are arranged within bigger parts of the same structure. This concept of a cascading organization seems ideally suited to chromatin organization, where groups of smaller domains come together to form larger structures, which again come together to form yet larger structures, and so forth (1, 4–6, 8, 11–15). Thus, while we here addressed large-scale euchromatin organization as a specific example, cascade processes in our opinion have potential also for the description of other types of chromatin and other length scales of organization.

A typical zebrafish embryonic nucleus is ≈ 10 *μ* m in diameter, while our imaging approach can resolve euchromatin organization down to 100 nm or finer. Our investigation thus covers at least two orders of magnitudes, at the level of large-scale euchromatin organization. Technical advances in the recent years, specifically in localization microscopy and electron microscopy tomography, have made available yet finer scales of chromatin organization (4, 5, 7–9). The view of a cascade process, which describes how chromatin domains at smaller levels are hierarchically organized into domains at the next larger levels, might offer a useful language to make sense of the structures revealed by these new techniques. An interesting question in this context would be whether the same cascade parameters apply across scales, or if the parameters change over the different scales of chromatin organization.

We would like to point out that the multiplicative cascade description of euchromatin packing simplifies the information contained in a full microscopy image to only four numbers. While this set of numbers is merely a phenomenological description of the domain-in-domain organization of euchromatin, these numbers can be used to decide whether hypothetical mechanisms of euchromatin organization are credible. A hypothetical mechanism that leads to a cascading process with the same parameters as inferred from microscopy images is credible, a process that does not result in the same parameters may be classified as not in line with the data. Such an assessment is distinctly simpler than the comparison against full image data. Considering that microphase separation has been proposed as a mechanism for the organization of different chromatin types and chromatin in general (3, 31–35), such a testing approach that is based in concrete experimental data seems relevant.

The description of euchromatin organization in terms of multiplicative cascades provides a fresh opportunity to test models of molecular access to specific targets. The question of how fast and with what probability for example transcription factors can reach their genomic target sites has been addressed previously (36). In this context, it was recognized that relatively inaccessible chromatin and the relatively more accessible nuclear space form an interspersed fractal organization, which would directly impact exploration of the intra-nuclear space by various macromolecules (37). Similarly, multiplicative cascades have previously been used to answer related questions, such as how pressure-injected fluids permeate polymeric networks (38). The use of multiplicative cascades would allow an abstract treatment of such questions. On the one hand, idealized mesh-works generated from cascade processes can be studied with more theoretical rigor and efficiency than real data. On the other hand, predictions from such theoretical work can, based on our fitting to microscopy images, be directly related to specific patterns of euchromatin organization.

Multiplicative cascades produce self-similar structures, which appear the same independently of the length scale they are viewed at. Such structures are also often described as fractals. The concept of fractal organization has also been applied successfully to chromatin organization, as studied by microscopy (39, 40), or x-ray scattering experiments (41). In these and other previous works, the fractal dimension or the strength of self-similarity was assessed. These analysis approaches can be used in combination with the multiplicative cascade approach we suggest here. However, in contrast to these previous approaches, multiplicative cascades do not only characterize experimental data, but can generate synthetic image data without further modifications or assumptions. Further, once the cascading rule is determined, theoretical models that underlie the observed self-similarity can be tested in a much simplified manner.

While our analysis supports the multi-scale, potentially scale-free organization of euchromatin, a microphase separation theory would imply the formation of patterns with characteristic length scales. The question arises how such a theory could still result in the formation of scale-free patterns. One proposed mechanism for microphase separation is the block copolymer nature of the genome, with subregions of different affinities (3, 31). If the subregions of such block copolymers would exhibit a scale-free organization in linear space, a scale-free organization in the 3D space filled by such polymers can be intuitively expected. Such scenarios further raise the question how such a scale-free organization of the block copolymers could be brought about. Here, recent theoretical work suggests that local, catalytically self-amplifying chromatin marks can establish microphase patterns (33, 35). Seeing that self-amplifying processes are frequently implied in the formation of scale-free patterns, such models seem promising candidates.

## CONCLUSIONS

The overall goal of this study was to capture the domain-within-domain packing of euchromatin in a process that describes not one particular length scale, but rather how the organization at one scale connects to the organization at other scales. We identified multiplicative cascades as an abstract process that can capture the multi-scale chromatin organization seen in 3D STED super-resolution microscopy images to a set of four parameters. When we applied this analysis to euchromatin organization in pluripotent zebrafish embryos, we found that different types of euchromatin organization systematically map to different parameter combinations. In particular, we find that when going from large to smaller scales, chromatin becomes more asymmetrically distributed to sub-regions in patterns that show stronger chromatin compaction. This asymmetry was most pronounced in euchromatin organization that can be explained by phase separation of the euchromatin polymeric chains. The asymmetry in the cascade parameters was still present, but not as pronounced in dispersed euchromatin organization patterns that are associated with a microemulsion or microphase-separated state. These particular sets of parameters provide a well-posed testing scenario for mechanistic models of euchromatin organzation, such as microphase separation via microemulsification (16).

## AUTHOR CONTRIBUTIONS

VZ, LH designed research. AN, LH performed experiments. VZ, HSK, LH analyzed data. LH wrote the manuscript

## ACKNOWLEDGMENTS

This work is funded via the Helmholtz Program Biointerfaces in Technology and Medicine (BIFTM); HSK, VZ – Volkswagen Foundation initiate “Life?”; VZ, LH – Deutsche Forschungsgemeinschaft priority program “Molecular Mechanisms of Functional Phase Separation” (DFG SPP-2191). Images were acquired at the Karlsruhe Center for Optics and Photonics (KCOP). The fluorescence dyes SiR-DNA, SPY650-DNA, and SPY595-DNA were provided free of charge by Spirochrome.

## SUPPLEMENTARY MATERIAL

**Figure 5:**
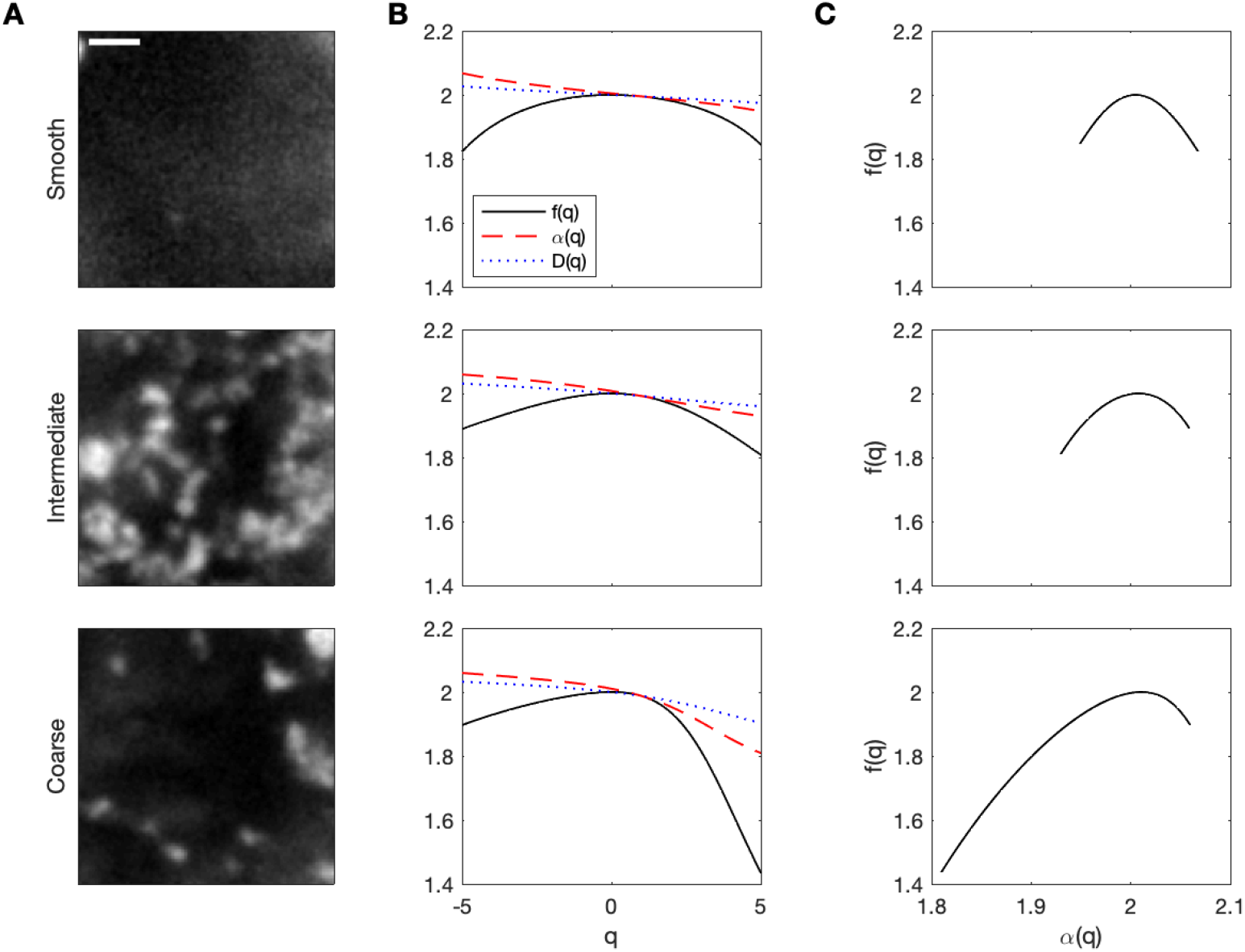
Euchromatin organization shows characteristics of multi-fractal organization. A) Example microscopy images that represent the different types of euchromatin organization seen in our data. The displayed images are the same as in Fig. 4 of the main article, and are used here for multi-fractal analysis. Scale bar: 1 *μ*m, note that a larger area of the stacks was used for analysis than in Fig. 4. B) Characteristic statistics *f* and *α* as well as the generalized dimension *D* are displayed for different analysis exponents *q*. C) *f* –*α* plots as commonly used to characterize the multi-fractal nature of a given system. A pronounced arc that spans a range of *α* values is considered indication of multi-fractal patterns. The displayed arcs span values that are similar to previous studies that concluded that systems exhibited multi-fractal patterns (21, 22, 28).

**Figure 6:**
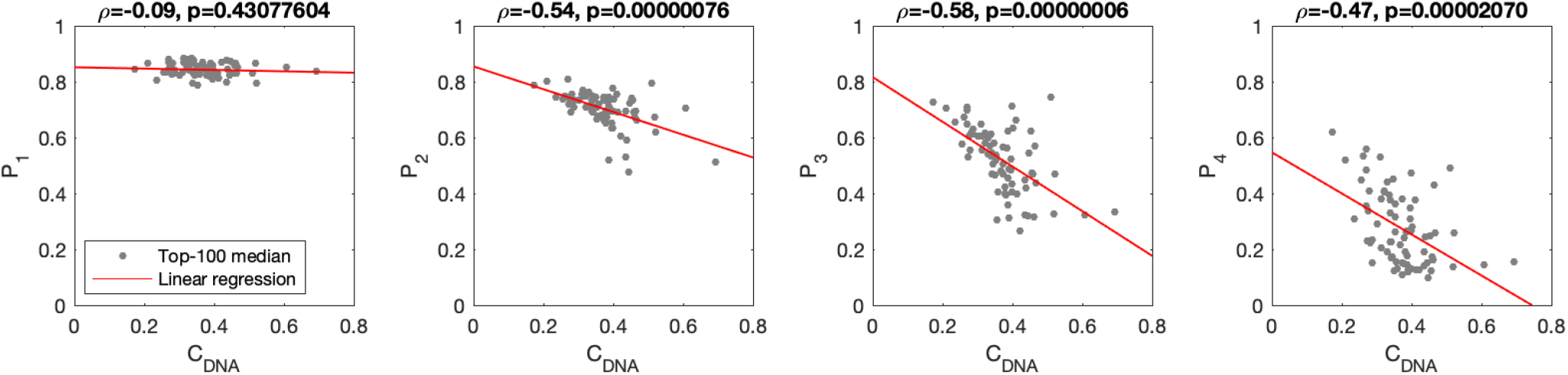
Higher contrast in DNA images correlates with more asymmetric multiplicative cascade parameters. Gray circles are medians of the top-100 parameter sets (evaluated based on variance, skewness, kurtosis comparison). Linear fits (red line) are shown. The Pearson correlation coefficient (*ρ*) and the p value for the particular values of *ρ* are indicated for each cascade parameter.

